# Vascular endothelial cadherin remodeling is associated with endothelial barrier dysfunction in chronic subdural hematoma

**DOI:** 10.64898/2026.09.27.754779

**Authors:** Kristine Ravina, Mark C. Renton, Jordan Darden, Clare Dennison, Eric A. Marvin, Scott R. Johnstone

## Abstract

**Background:** Chronic subdural hematoma (cSDH) remains a significant and increasingly prevalent cause of morbidity in older adults. Disease progression is sustained by de novo formed highly vascularized pseudomembranes and a persistent cycle of fluid accumulation and hematoma expansion. Although inflammatory and angiogenic pathways have been implicated in cSDH pathogenesis, the endothelial mechanisms underlying persistent vascular leak remain poorly understood. We investigated the cSDH vascular microenvironment in human patients and examined the role of vascular endothelial cadherin (VECAD) remodeling in cSDH endothelial dysfunction.

**Methods:** Human cSDH membranes and fluid were evaluated by multiplex profiling, histology, immunofluorescence, immunoprecipitation, western blotting and exploratory bulk RNA sequencing. A human brain endothelial cell (HBEC) line was exposed to patient-derived cSDH fluid to assess VECAD remodeling, endothelial barrier permeability, and proliferation.

**Results:** Multiplex profiling of patient-derived cSDH fluid detected mediators associated with endothelial cell (EC) activation, extracellular matrix remodeling, inflammation, and angiogenesis. In patient pseudomembranes, discontinuous VECAD coincided with erythrocyte extravasation. Two hyperdense (high-risk) specimens showed more extensive disruption than the hypodense (low-risk) specimen while vessel counts appeared similar. Bulk RNA sequencing identified an angiogenic environment and enrichment in cadherin regulation. A soluble VECAD (sVECAD) isoform was detected in cSDH fluid. Exposure of HBECs to patient-derived cSDH fluid increased EC proliferation, cytoplasmic VECAD (cVECAD) accumulation, and EC barrier permeability. Concurrent dexamethasone or minocycline treatment attenuated VECAD cleavage.

**Conclusions:** VECAD disruption is a prominent feature of cSDH and is associated with EC barrier dysfunction. These findings support further evaluation of soluble VECAD as a candidate biomarker and of junction-preserving pathways as potential therapeutic targets.

## Introduction

Chronic subdural hematoma (cSDH) is one of the most common neurosurgical conditions in older adults, with an increasing incidence associated with population aging and widespread antithrombotic medication use.^1-4^ Current management includes repeated surgical drainage, and adjunctive middle meningeal artery embolization which adds procedural risk and cost in a vulnerable patient population.^2,5^ The pathogenesis of cSDH involves a self-perpetuating cycle of inflammation, angiogenesis, and recurrent microhemorrhages resulting from disruption of fragile neovessels within the cSDH pseudomembrane.^6^ Although numerous inflammatory cytokines and angiogenic mediators have been implicated^7^, the mechanisms responsible for persistent endothelial cell (EC) barrier dysfunction remain incompletely understood.

Progressive cSDH enlargement is associated with continued leakage from immature pseudomembrane neovessels, suggesting that disruption of EC adherens junctions may contribute to vascular instability. Vascular endothelial cadherin (VE-cadherin; VECAD) is the primary adherens junction protein that maintains EC-EC cell adhesion and vascular barrier integrity.^8,9^ Loss or proteolytic cleavage of membrane-associated VECAD can increase EC permeability, increasing vascular leak. Proteolytic cleavage can degrade VECAD and release extracellular cadherin domains, producing soluble VECAD (sVECAD). Circulating sVECAD has emerged as a biomarker of endothelial injury in sepsis, acute kidney injury, vasculitis, aortic disease, and subarachnoid hemorrhage linked with inflammation.^8,10-14^ Whether VECAD remodeling is associated with EC barrier dysfunction in cSDH remains unknown.

We investigated VECAD in human cSDH using complementary patient-derived tissue and hematoma fluid, together with an *in vitro* human brain EC model. We examined VECAD localization in vascularized pseudomembranes, evaluated sVECAD in hematoma fluid, and determined the effects of patient-derived cSDH fluid on VECAD cleavage, EC barrier permeability, and proliferation in human brain EC in culture. Our findings identify VECAD remodeling as a prominent feature of cSDH-associated EC dysfunction and support further investigation of EC adherens junction disruption as a biomarker of vascular instability and a potential therapeutic target in cSDH.

## Methods

*See Supplement*

## Results

### The cSDH microenvironment contains mediators associated with vascular remodeling and inflammation

cSDH fluid from five patients were analyzed by multiplex human cytokine, chemokine, growth factor and metalloproteinase assays designed to characterize the local microenvironment. All analytes detected above the lower limit of quantification of the assay are displayed, revealing abundant expression of growth factors including PDGF-AB/BB, PDGF-AA, VEGF-A, and VEGF-AA, as well as immune factors including IL-6, IL-8, MCP-1, MMP-2, and IP-10 across all patient samples (**Fig. 1A, Supplemental Data 1**). Hallmark pathway analysis of detected analytes identified enrichment of angiogenesis, TNF-α signaling, and epithelial-to-mesenchymal transition (**Fig. 1B**). Histology and VECAD immunofluorescence of surgically removed pseudomembranes showed fibrous, cellular tissue with extensive vascularization, numerous hemorrhagic vascular spaces, and evidence of erythrocyte extravasation and inflammatory cell accumulation (**Fig. 1C**). Three patients were graded using the CT-based framework described by Stanišić and Pripp^15^ for cSDH phenotype: one hypodense/ low-risk and two predominantly hyperdense/ high-risk (**Fig 1D**). VECAD discontinuities spatially coincided with erythrocyte extravasation in representative regions, consistent with focal EC barrier disruption (**Fig 1D, arrows**). The two high-risk specimens showed more extensive VECAD discontinuity and increased extravasation than the low-risk patient, whereas vessel counts appear similar (**Fig 1D**).

**Figure 1.**
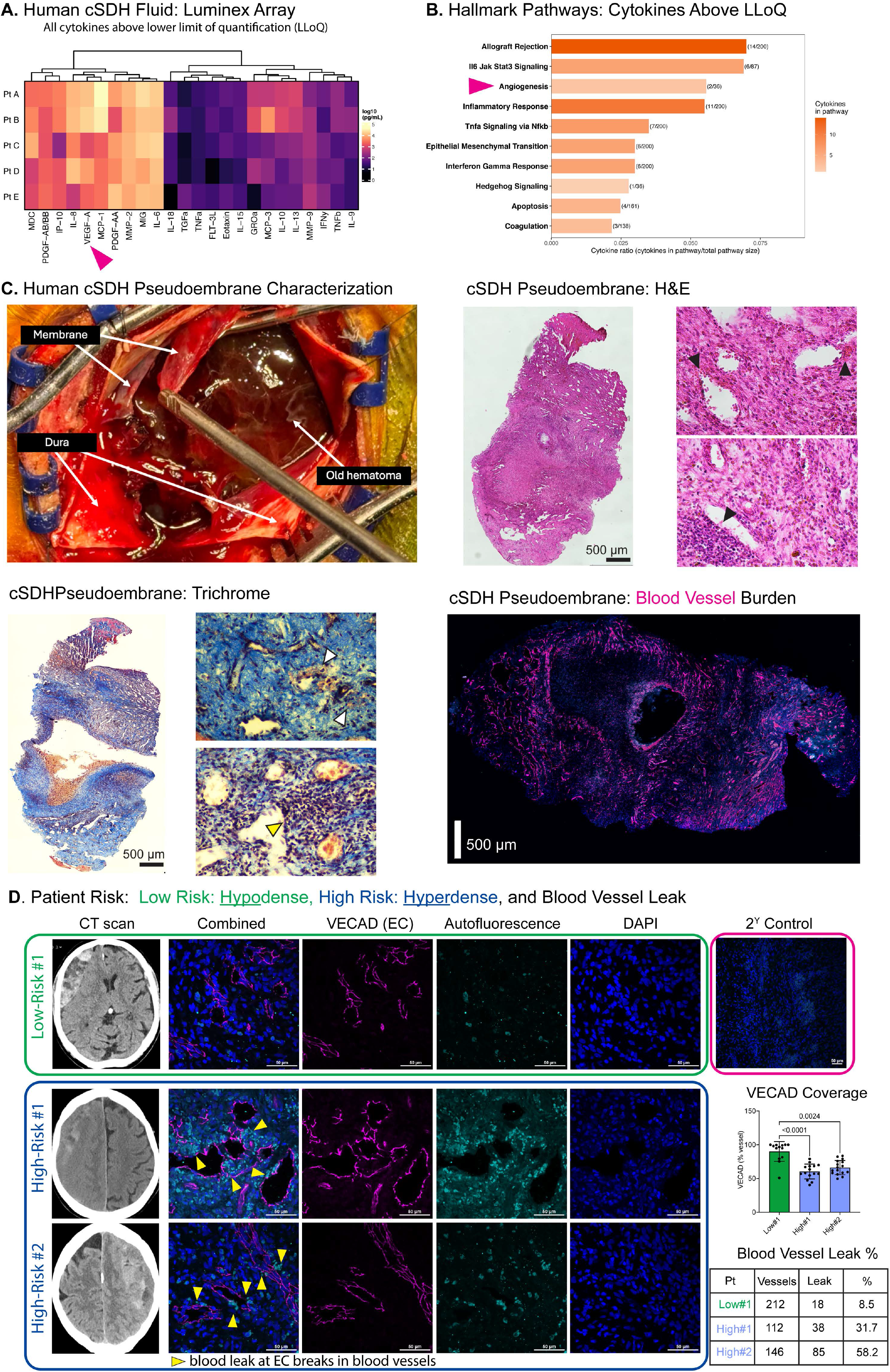
The cSDH microenvironment is characterized by inflammation, vascular remodeling, and EC barrier dysfunction. Multiplex profiling of cSDH fluid from five patients displaying clustered analytes detected above the lower limit of quantification (LLoQ) (**A**). Top 10 enriched Hallmark gene-set representation based on the measured analyte panel (**B**). Arrows highlight VEGF and angiogenesis (**A, B**) Location and histological features of the cSDH pseudomembranes, including hematoxylin and eosin and trichrome staining, colors are shown as blue/ green for collagen and connective tissue, red/pink for muscle fibers and cytoplasm, brown/red for red blood cells, and black/dark brown for cell nuclei (**C**). Immunofluorescence of VECAD (neovessels, EC) shows abundance of blood vessels in a pseudo-membrane (**C, lower right**). Head CT scans of patients with hypodense/ low-risk cSDH (n=1) and hyperdense/high-risk cSDH (n=2) (**D**). Immunofluorescence of VECAD neovessles (magenta) in low/ high-risk pseudomembranes, with quantification of vessel number and percentage of blood leak (green) at regions of VECAD breaks (arrows, **D**). n=13-16 vessels, Kruskal-Wallis multiple comparison test (**D**).

### cSDH pseudomembranes show angiogenic and junction-related transcriptional signatures

Bulk RNA sequencing compared patient pseudomembranes from three patients with matched peripheral blood RNA (n=3; low-risk hypodense n=1, high-risk hyperdense n=2, **Fig 2A**). Gene Ontology (GO) biological process (bp)_pathway analysis of differentially expressed genes (DEGs) in membrane compared to blood revealed enrichment of processes related to cell motility, cell proliferation, matrix organization, and angiogenesis (**Fig. 2B**). Patient-level heatmaps showed heterogeneous expression of junction-related genes including CDH5 and TJP1 (**Fig. 2C**). In vitro, cSDH fluid treatment significantly increased HBEC proliferation (**Fig. 2D**).

**Figure 2.**
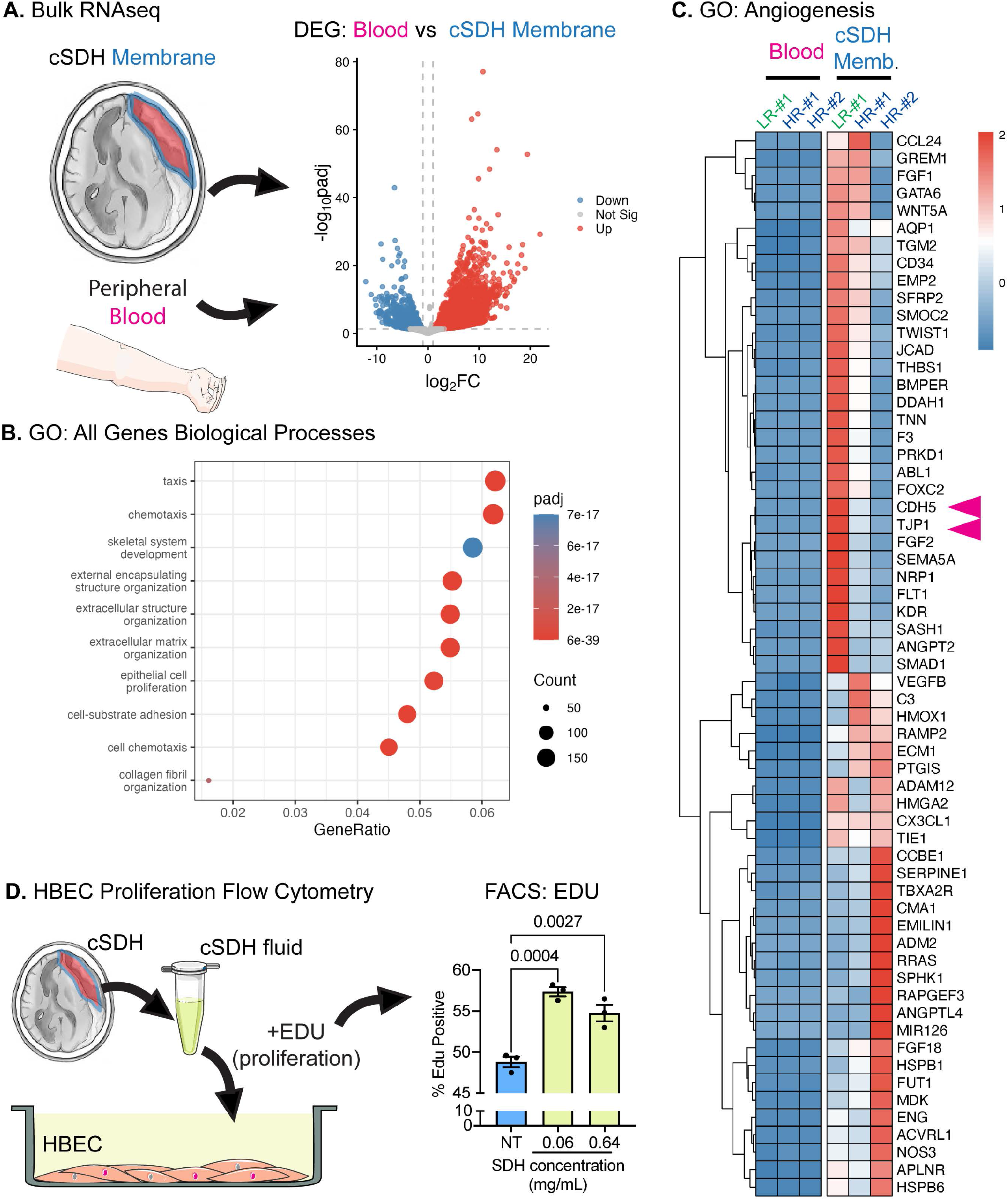
cSDH membranes have elevated markers of angiogenesis. Bulk RNA sequencing comparing pseudomembranes with matched peripheral blood from three patients with volcano plot of ‘DESeq2’ differentially expressed genes (DEGs) identified with |log_2_(Fold Change (FC))| > 1 and an adjusted p-value (padj) < 0.05 (**A**). Top 10 Gene Ontology (GO) biological process (bp) upregulated pathways (cSDH membrane vs blood) ranked by gene ratio (**B**). Patient-level heatmap ‘DESeq2’ normalized gene expression z-scores for the enriched Angiogenesis term (GO:0001525) for low-risk (LR#1, n=1) and high-risk (HR#1/HR#2, n=2), including CDH5 and TJP1 (**arrows, C**). 5-ethynyl-2’-deoxyuridine (EDU) proliferation flow cytometry in HBEC with no treatment (NT) or cSDH fluid treated (0.06 mg/mL and 0.64 mg/mL) (**D**). Percentage EDU positive cells, n=3, one-way ANOVA, and Dunnett’s multiple comparison test across groups.

### Patient cSDH fluid promotes VECAD breakdown and EC barrier dysfunction in vitro

KEGG pathway analysis of membrane-versus-blood DEGs identified enrichment of junction-related pathways including “Cadherin Signaling” (**Fig 3A**). VECAD is the primary EC cadherin and regulates junctional integrity. VECAD is a 120-125 kDa membrane-bound protein. Proteolytic processing can generate an extracellular soluble form containing cleaved extracellular VECAD repeats (sVECAD, ∼90kDa) which is associated with disruption of barrier function,^16^ and a cytoplasmic endosomal c-terminal cleaved VECAD (cVECAD, ∼95kDa) resulting from degradation.^17^ Immunoprecipitation and western blotting of clarified cSDH fluid detected an ∼90 kDa extracellular-domain VECAD compared to the 125 kDa VECAD in cultured EC, indicative of sVECAD presence in cSDH fluid (**Fig. 3B**). Exposure of HBEC to cSDH fluid from five individual patients (60 μg/mL for 24 hr) was associated with expression of an intracellular ∼95kDa VECAD isoform, indicative of cVECAD (**Fig. 3C**). In Transwell assays cSDH fluid exposure significantly increased HBEC permeability to 4 kDa FITC dextran (**Fig. 3D**). Concurrent treatment of HBEC with cSDH fluid and anti-inflammatory/anti-angiogenic drugs minocycline and dexamethasone reduced the presence of cVECAD in cSDH treated HBEC (**Fig. 3E)**

**Figure 3.**
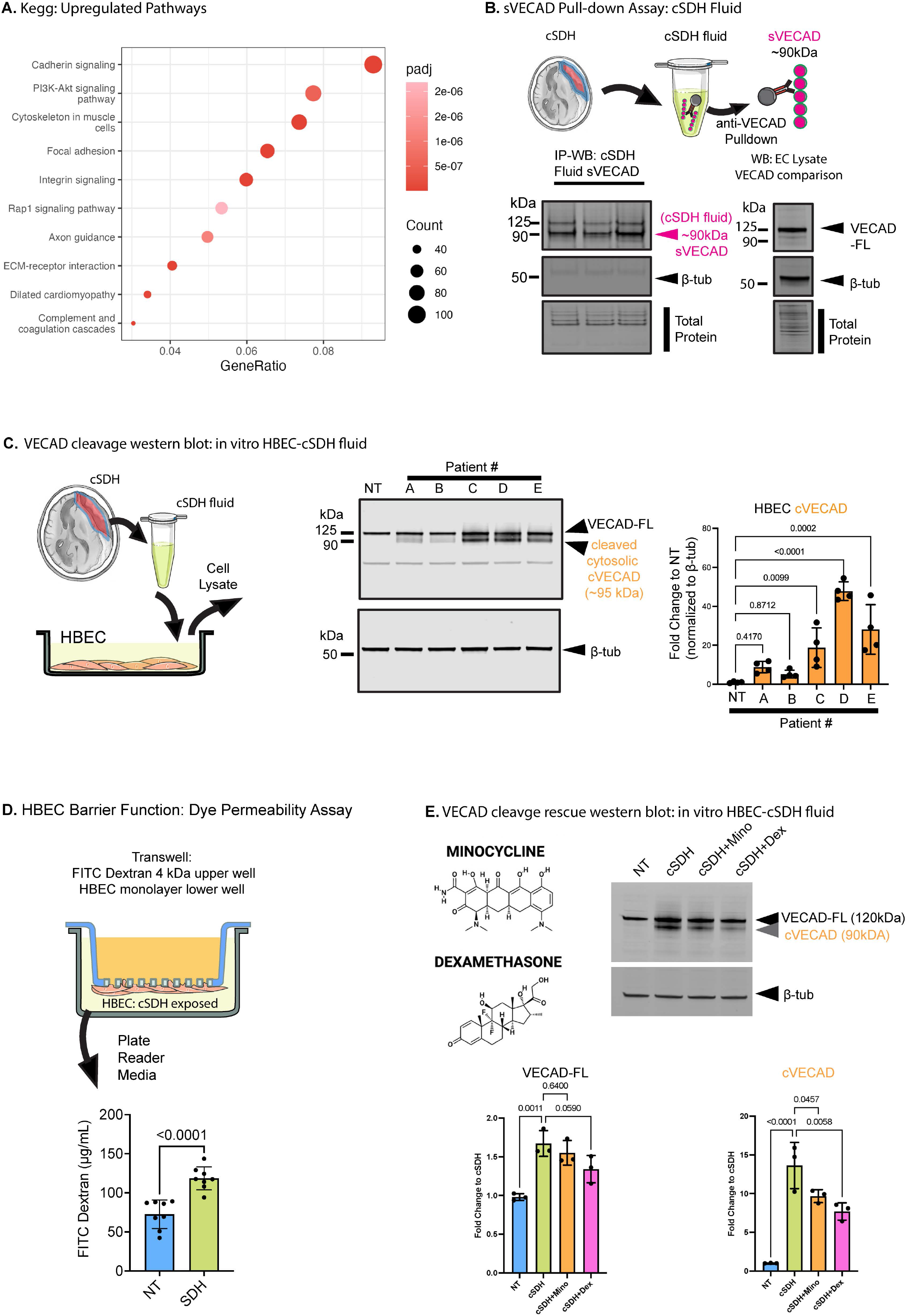
*Patient cSDH fluid contains sVECAD and promotes* VECAD cleavage associated with increased HBEC permeability. Top 10 KEGG upregulated pathways (cSDH membrane vs blood) ranked by gene ratio (**A**). Immunoprecipitation-western blot (IP-WB) of patient cSDH fluid, with detection of sVECAD (∼90 kDa), and western blot of VECAD-full length (VECAD-FL, ∼125 kDa) in untreated HBEC (**B**). HBEC western blot following treatment with patient cSDH fluid using anti-VECAD showing full length (VECAD-FL, ∼125 kDa) and cleaved VECAD (cVECAD, ∼95kDa) (**C**). Quantification of cleaved VECAD, n=5 patient treatments, n=4 replicates, normalized to β-tubulin, one-way ANOVA, and Dunnett’s multiple comparison test across groups. Transwell dye leak assay in HBEC treated with cSDH fluid for 24 hr (D). n=8, unpaired t test. Western blot of HBEC treated with cSDH concurrent with minocycline or dexamethasone (**E**). Quantification of VECAD-FL and cVECAD, n=3 replicates, normalized to β-tubulin, one-way ANOVA, and Dunnett’s multiple comparison test across groups.

## Discussion

In this exploratory study of human cSDH, VECAD discontinuity was observed in patient pseudomembrane vessels coinciding with erythrocyte extravasation, which appears to be increased in high-risk patients. cSDH fluid contained mediators associated with inflammatory and vascular-remodeling and we identified an approximately 90 kDa VECAD fragment that is consistent with previous reports for sVECAD. In vitro, exposure of HBEC to cSDH increased proliferation and permeability associated with increases in VECAD breakdown, that was attenuated by minocycline or dexamethasone treatment. Together these findings identify adherens junction remodeling as a candidate mechanism and pharmacologically targetable component of cSDH vascular dysfunction.

Persistent or recurrent cSDH is associated with continued fluid leakage from immature neovessels.^6,7^ Prior work has emphasized inflammatory cytokines, growth factors, and angiogenic mediators involved in pseudomembrane formation and vascular instability.^7,18,19^ Consistent with this literature, multiplex profiling demonstrated that cSDH fluid contains a complex milieu of proteins associated with EC activation, extracellular matrix remodeling, inflammation, and angiogenesis. Our data demonstrate that cSDH fluids contain factors associated with EC growth including VEGF-A, and cSDH fluid exposure increased EC proliferation in vitro. These results support a proangiogenic environment that promotes formation of immature blood vessels that are prone to leak. This presents as a potential pharmacologic target to limit vascular dysfunction. Targeting VEGF-A (Bevacizumab) has shown promise in preclinical and early clinical studies.^20-22^ Yet, there are concerns of side effects, including hypertension, hemorrhage and proteinuria.^18,23^ Additionally, VEGF levels are elevated in primary SDH, but are not no significantly different in cSDH recurrence, suggesting that VEGF may not be the primary factor leading to recurrence.^24^ We observed increases in other growth factors including PDGF-AB/BB, PDGF-AA which promote cell proliferation but are also associated with recruitment of pericytes, which are involved in vessel stability^25-27^. Despite increasing characterization of the soluble cSDH microenvironment, our study does not identify a single causal mediator. The downstream EC mechanisms responsible for persistent vascular leak remain comparatively underexplored and further studies are warranted to understand the vascular dysfunction in cSDH.

CT morphology has been associated with recurrence risk in clinical cohorts. Predominantly hypodense hematomas exhibit fewer vascularized pseudomembranes and lower recurrence rates than hyperdense hematomas, which are characterized by abundant pseudomembranes and a higher recurrence risk.^15^ The three pseudomembranes used in this study comprised two predominantly hyperdense specimens and one predominantly hypodense specimen. Our study did not record recurrence. Although all three pseudomembranes were vascularized with similar vessel counts, the hyperdense specimens showed more extensive VECAD discontinuity and erythrocyte extravasation than the hypodense specimen. This suggests that factors regulating vessel stability, rather than pure vessel growth are involved. As such, EC barrier disruption may contribute to the heterogeneous biological behavior of these lesions. EC barrier function is governed by coordinated regulation of junctional complexes, where VECAD, a canonical adherens junction protein, stabilizes cell-cell contacts and restrains paracellular permeability.^9^ VECAD restricts the release of free plasma proteins from blood into tissue.^28^ Disruption of VECAD organization can cause barrier failure, promoting blood/plasma leak and increasing paracellular permeability.^8,9,12^. While our data suggest that EC barrier failure is a significant factor, our findings do not currently establish the effect of CT phenotype or predict recurrence. Further evaluation of VECAD organization and EC junction preservation in a larger clinical cohort is needed to understand the importance of junctional disorganization as a contributor to ongoing vascular leakage.

At the plasma membrane, VECAD adhesive activity is supported by interactions with vascular endothelial protein tyrosine phosphatase (VE-PTP), and disruption of this complex can increase endothelial permeability.^29,30^ In inflammatory and vascular disorders, proteolytic ectodomain shedding releases fragments including sVECAD comprised of the five extracellular cadherin repeating domains (EC1-5) associated with VECAD-VECAD interactions, and VE-PTP binding.^8,10,31^ Clinical studies report circulating sVECAD increases in severe systemic inflammatory states with microvascular leakage.^8,10,13^ Circulating sVECAD is elevated and linked with adverse outcomes in sepsis, acute kidney injury, vasculitis, and aortic dissection.^11,12,14,32^ Recent mechanistic work establishes sVECAD as a causal mediator, not just a marker, with recombinant sVECAD inducing loss of barrier function in vitro and in vivo by perturbing VECAD:VE-PTP.^8^ sVECAD functions as a decoy that sequesters VE-PTP away from membrane VECAD, destabilizing junctions through loss of homotypic adhesion and aberrant cytoskeletal contraction. Our *in vitro* data using patient cSDH fluid show that cSDH fluid contains high levels of sVECAD and that cSDH fluid induces VECAD breakdown in cell culture model, evident by production of cVECAD and barrier leak. However, we did not directly test if sVECAD caused the effects of cSDH. Future studies should assess if sVECAD in cSDH fluid should be considered a biomarker and mediator of vascular instability.

Despite the rising burden, treatment approaches proposed to date including surgical interventions and pharmacotherapeutics, remain limited by unacceptable systemic side effects, high recurrence rates, high cost, complication risk or lack of evidence. We chose to investigate the effects of both dexamethasone and minocycline as both target pathways are linked with VECAD degradation including MMP, ADAM and VEGF pathways. Systemic corticosteroids such as dexamethasone have been clinically evaluated, but current data does not support their use due to the high risk of systemic complications and local drug delivery remains largely unexplored.^24,33^ In previous studies, treatment in patients did not reduce VEGF, or anti-inflammatory profile (IL-6, IL-8, and IL-1α), although infiltration to the subdural space may be a limiting factor in oral delivery^24^. In our in vitro studies, dexamethasone attenuated VECAD cleavage. Glucocorticoid signaling has been associated with upregulation of junctional proteins including VECAD, while downregulating proteases including MMP-9 which is associated with junctional breakdown, and VEGF which is linked to VECAD phosphorylation.^24,34,35^ Thus, dexamethasone effects may reflect changes to inflammatory signaling, junctional protein expression, protease activity or VEGF signaling rather than direct inhibition of VECAD cleavage. Minocycline also attenuated cVECAD. Minocycline is a well-characterized broad-spectrum metalloproteinase inhibitor that may cause its effects on MMP/ADAM-dependent cleavage of VECAD linked to further shedding of sVECAD.^36^ While this pathway may not directly produce cVECAD, it may limit barrier function degradation, resulting in less VECAD cleavage and reduced sVECAD released to the surrounding environment, limiting feed forward signaling. However, further studies are required to understand the mechanism controlling VECAD, including the actions on associated enzymes including calpain, as the fragments we identify are consistent with calpain-mediated cleavage of VECAD, endosomal uptake and VECAD disorganization.^17,37^

This study has several limitations. The patient cohort was small, which limits interpretation of the CT-phenotype and RNA sequencing comparisons. Larger patient cohorts are needed to determine whether VECAD cleavage predicts recurrence or clinical outcomes. The *in vitro* experiments used patient-derived cSDH fluid and cultured HBEC, which cannot fully recapitulate the complex microenvironment of cSDH. Regardless, VECAD cleavage was a consistent factor in treatment using cSDH fluid from multiple patients. Finally, while these data demonstrate VECAD cleavage and EC barrier dysfunction, the upstream proteolytic mechanisms responsible for VECAD processing remain undetermined and are under active investigation. While we have shown abundant sVECAD in cSDH fluid, multiple factors are present in the fluid that may have effect. Future studies are required to understand the exact environment surrounding cSDH pseudomembranes that promotes not only angiogenesis but also regulates vascular leak.

In summary, VECAD disruption in cSDH pseudomembranes is associated with EC barrier dysfunction. These findings support adherens-junction remodeling as a candidate component of cSDH vascular pathology and sVECAD as a candidate biomarker. Larger patient cohorts, and direct mechanistic experiments are required prior to risk prediction or junction-preserving therapies can be expected to improve clinical outcomes.

## Figure legends

**Supplemental Data 1**: Multiplex human cytokine, chemokine, growth factor and metalloproteinase assay data

## Supplemental methods

### Human Specimens

cSDH fluid and peripheral blood were collected from patients undergoing surgical evacuation of cSDH after obtaining the appropriate consent under Carilion Clinic institutional review board-approved tissue biorepository protocol (IRB# 20-1197). cSDH patient specimens were identified and retrieved for analysis under Carilion Clinic IRB 24-2173 cSDH proof-of-concept study. Only cSDH fluid was available from a total of five patients while cSDH membranes, cSDH fluid, and peripheral blood were available from three patients. Samples were de-identified before analysis. cSDH fluid was centrifuged to remove cellular debris, aliquoted, and stored at -80°C until analysis. Total protein was measured by bicinchoninic acid assay (ThermoFisher, #A55861). Pseudomembranes were divided into ∼1cm^2^ sections and either fixed in 4% PFA for histology or stored at -80°C for biochemical analyses and RNA sequencing. Blood was collected for RNA in PAXgene tubes (Qiagen #762165). Patient computed tomography scan images were used to radiographically classify low-risk (predominantly hypodense, n=1) or high-risk (predominantly hyperdense, n=2) cSDH based on the grading framework described by Stanišić and Pripp.^15^ Recurrence outcomes were not evaluated; therefore, these imaging categories were not treated as observed clinical-risk groups.

### Multiplex cytokine profiling

Cytokines, chemokines, and growth factors were quantified in cSDH fluid using a commercially available human magnetic bead-based multiplex immunoassay (MILLIPLEX Human cytokine/chemokine/growth factor Panel A, MilliporeSigma HCYTA-60K-PX48) on an xMAP INTELLIFLEX® platform (Luminex Corp) with normal human serum control according to manufacturer instructions. Curves were fitted using Belysa Immunoassay Curve Fitting Software (MilliporeSigma). Metalloproteinases were measured in the same patient samples using a separate human magnetic bead multiplex assay. Concentrations (pg/ml), limit of detection, and lower limit of quantification (LLOQ) were calculated for each protein from standard curves using the manufacturer’s software. For heatmap visualization, analytes detected above LLoQ in every sample were log_10_-transformed, subjected to unsupervised Euclidean hierarchical clustering, and visualized with ComplexHeatmap (v2.22.0) in R (v4.4.3). For pathway analysis, detected analytes were mapped to human Hallmark gene sets using msigdbr (v25.1.1). Enrichment was determined by ranking gene sets by the total number of detected analytes in a set divided by the total number of genes in that set, and the top 10 highest-ranked sets were visualized using ggplot2 (v4.0.2). Raw and processed data files are supplied in Supplementary Data 1.

### Immunofluorescence and histology

Patient cSDH pseudomembranes were processed for hematoxylin and eosin (H&E) and trichrome histology. Immunofluorescence was performed as previously described^38^ using an antibody against VECAD c-terminal domain (EC, Sigma #Vli37). Whole tissue images were acquired by large-area scanning and vascularized regions were imaged to evaluate continuity of junctional VECAD staining. Sites of VECAD disruption were compared with areas of erythrocyte extravasation. Representative high-risk and low-risk specimens were selected according to the cSDH CT classification.^15^ Histological images were acquired on a Nikon Ci-L microscope, and immunofluorescent images were captured using Nikon NIE confocal microscope. Single images, maximum intensity Z-stack projection, along with tile scanning, were used to visualize samples. All quantitative analysis (vessel number and RBC extravasation) was performed using NIS Elements Advanced Research software (Nikon) by a researcher blinded to the sample classification.

### Human Brain EC Culture

Human cerebral microvascular endothelial cell line hCMEC/D3, (HBEC, Millipore, #SCC066) was maintained in vascular media (ATCC, #PCS-100-030) supplemented with endothelial cell growth kit (ATCC, #PCS-100-041) containing 2% FBS. Before experiments, cells were maintained in cell media containing low-serum (0.1%) containing no additional growth factors for three days. Cells were exposed to patient-derived cSDH fluid normalized to total protein content for the stated times and concentrations. Dexamethasone and minocycline were added concurrent with cSDH fluid.

### HBEC Permeability Assay

Transwell inserts (0.4 μm pore, Costar #3452) were coated with collagen (PureCol, Advanced Biomarix) and HBEC cultured to confluence on the underside and maintained in low serum media for three days. After exposure to pooled cSDH fluid (1 mg/mL, 24 hr), HBEC barrier permeability was assessed by adding 4-kDa FITC-dextran (1 mg/mL, Sigma #46944) added to the upper chamber. After 2 hr, fluorescence signal in the lower chamber media was quantified against a FITC dextran standard curve using a plate reader (Clariostar, BMG Labtech). Untreated cells (NT) and cell-free Transwells were used as controls.

### HBEC Proliferation Assay

HBEC were maintained for three days in low-serum media and then exposed for 24 hr in low serum media containing patient cSDH fluid (0.6 or 0.06 mg/mL total protein) and 5-ethynyl-2′-deoxyuridine (EdU, Invitrogen #C10634). At 24 hr cells were fixed in 70% ethanol and processed according to the manufacturer’s protocol. EDU incorporation indicating proliferative cells, was measured by flow cytometry (Sony SH800) and the percentage of EdU-positive cells calculated.

### Western Blotting and Immunoprecipitation

HBEC were processed for western blotting as previously described^39,40^. Soluble vascular endothelial cadherin (sVECAD) was immunoprecipitated using anti-VECAD antibody (R&D Systems #AF938, human VE-Cadherin Asp48-Gln593) cross-linked to magnetic beads (Sigma, Dynabeads #11201D). Immunoprecipitated proteins were detected by western blotting using antibodies recognizing the extracellular domain of VECAD (R&D Systems #AF938) and secondary fluorescent antibodies (Licor #926-32210) and normalized to β-tubulin (developmental studies hybridoma bank, #E7) or total protein (Licor, # 926-10011). Membranes were imaged (Licor Odyssey) and quantified (Image Studio 5.2.5, Licor).

### Bulk RNA sequencing

RNA isolated from three patients (two high-risk and one low-risk) pseudomembranes and peripheral blood were processed for bulk RNA sequencing. Pseudomembrane tissues (∼1cm^2^) were lysed in 350 µl of RLT lysis buffer and RNA was extracted (RNeasy Fibrous Tissue Mini Kit, Qiagen #74704) according to the manufacturer’s instructions, including the optional DNA removal step. RNA from matched patient blood stored in PAXgene tubes was extracted (PAXgene Blood RNA Kit V2, Qiagen #762164) according to the manufacturer’s instructions. Extracted RNA was quantified using a NanoDrop One spectrophotometer (ThermoFisher, USA). Aliquots of RNA (1.5 µg) were then diluted to equal concentrations (50 ng/µl) in RNase-free water. RNA integrity analysis, library preparation, sequencing, raw read count generation, and differential gene expression analysis were performed commercially by Novogene (Sacremento, CA, USA) using their Human mRNA Sequencing pipeline. Full parameters, protocols, and quality control reports are available on request, but are explained briefly below:

RNA integrity was assessed using the 5400 Fragment Analyzer System (Agilent Technologies, CA, USA). For membrane samples, mRNA library preparation was performed with poly A enrichment, and blood samples followed the GLOBINclear and non-directional mRNA library preparation pipeline. Library quality was verified using Qubit assay for quantification, Agilent Bioanalyzer for size distribution, and qPCR for molarity using adaptor-specific primers. Sequencing was performed on the NovaSeq X Plus (Illumina, USA) platform with 150 bp paired-end reads and a minimum of 6 G raw data. Raw sequencing data in FASTQ format were cleaned using custom in-house (Novogene) Perl scripts. Reads with adapter contamination, reads containing more than 10% uncertain nucleotides, and reads in which more than 50% of bases had a quality (Q) score below 5 were removed. Reads were then aligned to the human genome (hg38) using ‘HISAT2’ (v2.2.1). Raw read counts were generated using ‘featureCounts’ (v2.1.1). For all downstream analysis raw count normalization was performed using ‘DESeq2’ (v1.46.0) in the R statistical analysis program (v4.4.1). To exclude low count genes prior to ‘DESeq2’ normalization, only genes with at least 10 counts in one sample were kept. Gene symbols were extracted based on provided Ensembl ID using ‘clusterProfiler’ (v4.14.6) and ‘org.Hs.eg.db’ (v3.20.0). ‘DESeq2’ normalized counts were log2 transformed to ensure a normal distribution. Euclidean distances were then calculated, and principal component analysis (PCA) was performed using ‘dist()’ and ‘prcomp()’, respectively, in base R.

Differential gene expression (membrane vs blood) was performed using ‘DESeq2’ with Benjamini-Hochberg multiple comparison adjustment. An adjusted p-value (padj) < 0.05 and an |log2(FC)| > 1 was set as the threshold for differentially expressed genes (DEGs). Volcano plots displaying -log10(padj) and log2(FC) were generated in using ‘ggplot2’ (v4.0.2).

Pathway enrichment analysis was performed on upregulated (log2(FC) > 1) DEGs with ‘clusterProfiler’ using the Gene Ontology (GO) biological process (bp) and Kyoto Encyclopedia of Genes and Genomes (KEGG) databases. The list of detected genes after low gene count exclusion was used as the background gene set. Pathways with a padj < 0.05 following Benjamini-Hochberg multiple comparison adjustment were considered statistically significantly enriched. The top 10 enriched pathways were plotted using ‘ggplot2’. An individual sample heatmap was generated by extracting ‘DESeq2’ normalized counts for all DEGs in the enriched angiogenesis GObp pathway and calculating a gene expression z-score for each sample. Data were then visualized using ‘pheatmap’ (v1.0.13).

Raw and processed data files have been deposited in the National Center for Biotechnology Information Gene Expression Omnibus under the accession number GSE348372.

### Statistical Analysis

Experiments were performed using biological replicates derived from independent patient samples where indicated. Data are presented as mean ± SD. Analyses were performed in Graphpad Prism 10.4.3.2, unless otherwise stated. Samples were assessed for normal distribution using a Shapiro-Wilk test, and outliers identified using the Prism robust regression and outlier removal (ROUT) test (Q =1%). 1-way multiple comparison ANOVA followed by Dunnett post-test was used for comparisons between >2 groups. The unpaired Student’s *t* tests were used for comparisons of 2 treatment groups. Kruskal-Wallis non-parametric tests were used for groups that did not display normal distribution. A minimum of n=3 was used for all statistical analyses, and n values for each experiment are stated in the figure legends. P values are shown for all statistical analyses, with a P value of less than 0.05 considered significant. Data represent mean ± SD.

